# Predicting provenance of forensic soil samples: soil DNA predicts habitat and environmental properties

**DOI:** 10.1101/390930

**Authors:** Camilla Fløjgaard, Tobias Guldberg Frøslev, Ane Kirstine Brunbjerg, Hans Henrik Bruun, Jesper Moeslund, Anders Johannes Hansen, Rasmus Ejrnæs

## Abstract

Environmental DNA is increasingly applied in ecological studies, including forensic ecology where eDNA from soil can be used to pair samples or reveal sample provenance. We collected soil eDNA samples as part of a large national biodiversity research project across 130 sites in Denmark. We investigated the potential for soil eDNA in predicting provenance in terms of environmental conditions, habitat characteristics and geographic regions. We used linear regression for predicting environmental gradients of light, moisture, soil pH and nutrients (represented by Ellenberg Indicator Values, EIVs) and quadratic discriminant analysis (QDA) to predict habitat class and geographic region. We found high predictive power for environmental gradients (R^2^ > 0.73). The discriminatory power of QDA in predicting habitat characteristics varied from high accuracy in predicting certain forest types, less accurate prediction of heathland and poor accuracy for geographic region. We demonstrate the application of provenance prediction in forensic science by evaluating and discussing two mock crime scenes. Here, we supplement with plant species lists from annotated sequences. Where predictions of environmental gradients and habitat classes give an overall accurate description of a crime scene, care should be taken when interpreting annotated sequences, e.g. due to erroneous assignments in GenBank. The outlined approach clearly demonstrates that basic ecological information that can be extracted from soil eDNA, contributing to the range of potential applications of eDNA in forensic ecology.

## Introduction

In ecological studies, bioindication is routinely used to infer environmental conditions and ecosystem properties, and to classify vegetation types (1–3). The link between species and environmental conditions is also the basis of the application of ecology in forensic science (4). In a wide range of disciplines – such as palynology, botany and entomology – pollen, plant fragments or insect remains are interpreted or analyzed by experts to impart ecological information to investigations (5–8). Similarly, forensic geoscience builds on the geological disciplines of inorganic soil analysis, i.e., soil classification, mineralogy, soil chemistry and physics (9).

Soil is commonly used as trace evidence in criminal cases to match two soil samples and thereby establishing a link between a suspect and a crime scene. In an investigative process, where for example the crime scene is unknown, soil trace evidence can also give valuable information on geographic origin or provenance and help narrowing a search. However, inorganic soil properties tend to vary at regional scales, which limits the precision of soil sample provenance (10), but recent digital signatures from x-ray fluorescence have demonstrated a potential for high precision prediction accuracy for local scale soil provenance (11).

Soil contains traces of DNA from the biota living in or above the soil. Plants are rooted in the soil, fungal mycelium grow through the soil and animals live, dwell or root through the soil, all leaving DNA traces behind. Even organisms above the soil eventually leave DNA in the soil as they defecate, exude secretes or die and decay. Extracting DNA from soil has been in use for decades in microbiology for characterization of bacterial communities in sediments (12). Also, microbial profiles from forensic soil DNA analysis have been used as a fingerprinting tool in forensic investigations (13, 14). With the development of metabarcoding of soil DNA, or environmental DNA from soil samples (from here onwards: soil eDNA; 15), high-throughput sequencing of marker genes allow simultaneous analyses of multiple species in a sample. These methods have rapidly found application in ecology (16), where DNA profiles are used in ecological studies and conservation to identify communities or quantify biodiversity (17, 18). Recently, these methods have also been employed in forensic ecology, where e.g., a “biological signature” of annotated taxa in soil DNA has been used to describe vegetation characteristics (19) and “fingerprints” of sequence composition at sites have been used to match forensic soil samples (20).

Here, we test the predictive power of soil eDNA with regard to provenance of an unknown soil sample, in terms of geography, abiotic conditions and habitat classes. More specifically, we address the questions: Can we use soil eDNA to predict the sample’s origin along environmental gradients in light, soil pH, nutrient status and moisture? Can eDNA predict common habitat classes, such as forest, heathland, rotational field? Can eDNA predict geographic origin? We used linear models to predict environmental gradients of focal samples. For habitat classes, we used quadratic discriminants analysis. Lastly, we explored to what degree expert evaluation of plant sequences could enhance the model predictions.

## Methods

### Sample sites

This study is based on data from the project “Biowide”, a large nationwide survey of biodiversity in Denmark, where multitaxon biodiversity and biodiversity drivers were investigated within the ecospace framework (21). We selected 130 study sites (40 m × 40 m) evenly distributed across five geographic regions in Denmark. Within each region, sites were placed in three clusters for logistic reasons, but with a minimum distance of 500 m. Site selection was stratified according to primary environmental gradients. We allocated 30 sites to cultivated habitats and 100 sites to natural habitats. The cultivated subset was stratified according to major land-use types and the natural subset was stratified according to gradients in soil fertility, soil moisture and successional stage. We deliberately excluded saline and fully aquatic habitats, but included temporarily inundated depressions as well as wet mires and fens. The final set of 24 environmental strata consisted of the following six cultivated habitat types: Three types of fields (rotational, grass leys, set aside) and three types of plantations (beech, oak, spruce). The remaining 18 strata were natural habitats, constituting all factorial combinations of: Fertile and infertile; dry, moist and wet; open, tall herb/scrub and forest. These 24 strata were replicated in each of the five geographical regions. We further included a subset of 10 perceived hotspots for biodiversity in Denmark, selected subjectively by public voting among active naturalists in the Danish conservation and management societies, but restricted so that each region held two hotspots. See Brunbjerg, Bruun (22) for a thorough description of site selection and stratification. From the 130 sites, we randomly chose 20 sites and from those chose two sites (one forested and one open habitat) to represent mock crime scenes for which we will present and discuss the soil provenance predictions in depth. The sites were located both on privately owned and public land. Oral permission was obtained from all landowners. All field work and sampling was conducted in accordance with Responsible Research at Aarhus University and Danish law.

Plant species nomenclature follows the database https://allearter.dk/.

### Soil eDNA metabarcoding

We collected soil from all sites and subjected it to metabarcoding through DNA extraction, PCR amplification of genetic marker regions (DNA barcoding regions) and massive parallel sequencing on the Illumina platform as described in Brunbjerg, Bruun (22). The soil sampling scheme included the mixing of 81 soil cores from each site in an attempt to get a representative sample. For this study, we used sequencing data from genes amplified with primers targeting eukaryotes, fungi, plants and insects. For eukaryotes we amplified part of the 18S region with primers 18S_allshorts (23, 24) with a slight modification of the forward primer (TTTGTCTGGTTAATTCCG). For fungi we amplified the ITS2 region with primers gITS7 (25) and ITS4 (26). For plants we amplified the ITS2 region with S2F (27) and ITS4. For insects, we amplified part of the 16S region with primers Ins_F and Ins_R (28).

The bioinformatic processing of the sequence data followed the strategy outlined in (22) based on the identification of exact sequence variants (29) by using the DADA2 pipeline (30) including post-clustering curation with the LULU algorithm (31) for the fungi, plants and insects in order to obtain reliable alpha diversity estimates for OTU data. OTUs that could not be assigned to the target group were removed from the datasets. Taxonomic affiliation was assessed by using the most widely applied name among the top hits (within 0.5%) from blastn against GenBank.

For insects and eukaryotes it was only possible to sequence 128 and 129 of the 130 samples, respectively.

### Predicting environmental gradients

To obtain meaningful predictors from OTU diversity, we used the first four ordination axes obtained from NMS-ordination (function metaMDS in package vegan, 32, default settings) of the four OTU community datasets, using abundance data (number of sequences per OTU) with square root transformation followed by Wisconsin double standardization. As response variables, we used environmental gradients represented by community mean Ellenberg Indicator Values (EIV, 33). These were obtained by averaging plant indicator values for the plant species present at the site, an approach commonly used in vegetation studies to assess local environmental conditions (e.g., 34). The validity of plant-based bioindication has been confirmed by direct measurement of the environmental conditions and by plant growth experiments (e.g., 35, 36). We used the EIV for soil fertility/nutrients, soil moisture, pH and light conditions (EIV N, M, R and L), which have been shown to be very useful for describing the variation in this dataset (see Fig. 3 in 22).

We used linear models to evaluate the relationship between OTU composition and environmental gradients (see Appendix S1). Model selection was based on AIC using the function stepAIC and backwards selection in package MASS (37, see Table S1). Individual models were constructed for each set of NMS axes based on the different primers and normality and heterogeneity were assessed by visual inspection of qq-plots, residual plots and histograms of residuals and indicated no problems with the final model. The best set of NMS axes was selected to predict EIV and 95% CI using leave-one-out cross validation. All analyses were performed in R-3.4.2 (38).

While most trained botanists and ecologists may be familiar with EIV, to most people they are hard to interpret in terms of meaningful vegetation types of environmental conditions. We have therefore aided interpretation by graphing the location of common vegetation types along the four Ellenberg gradients used in this study (Fig 1).

**Fig. 1.**
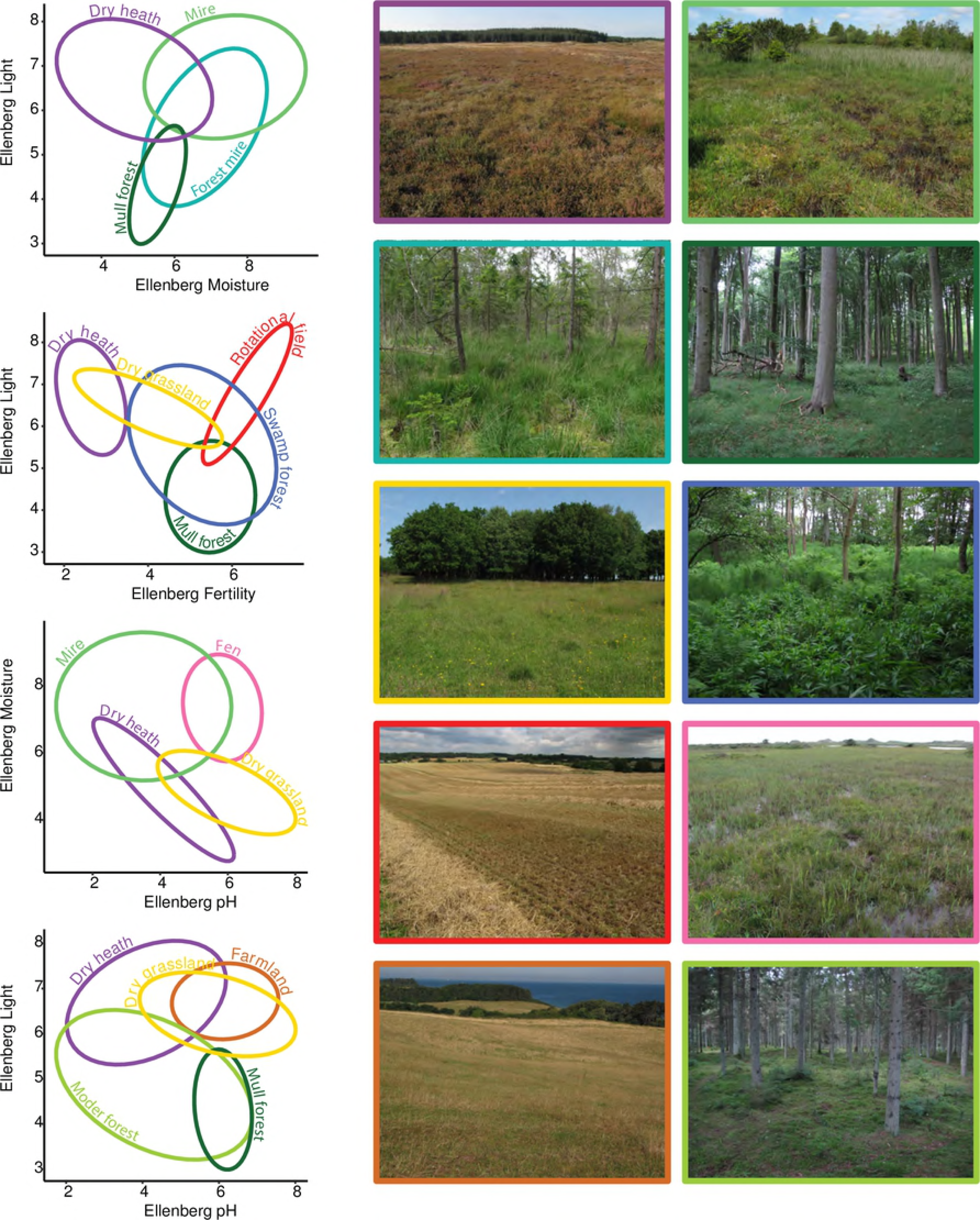
Interpreting Ellenberg Indicator Values. Ellipses showing the multivariate normal distribution of vegetation types plotted along Ellenberg indicator values for moisture, pH, light and nutrient status. Agricultural includes rotational fields, old fields and lays. The color of the ellipses correspond to characteristic sites depicted on the right.

### Predicting binary habitat classes

Another aim was to describe habitat classes typical of Denmark from the OTU composition in soil samples. We selected classes that are familiar to most people, easy to identify by non-ecologists and possible to recognize from a distance or from publicly available maps and orthophotos with some training (Table 1). The habitat classes were recorded as binary variables for each site.

**Table 1.**
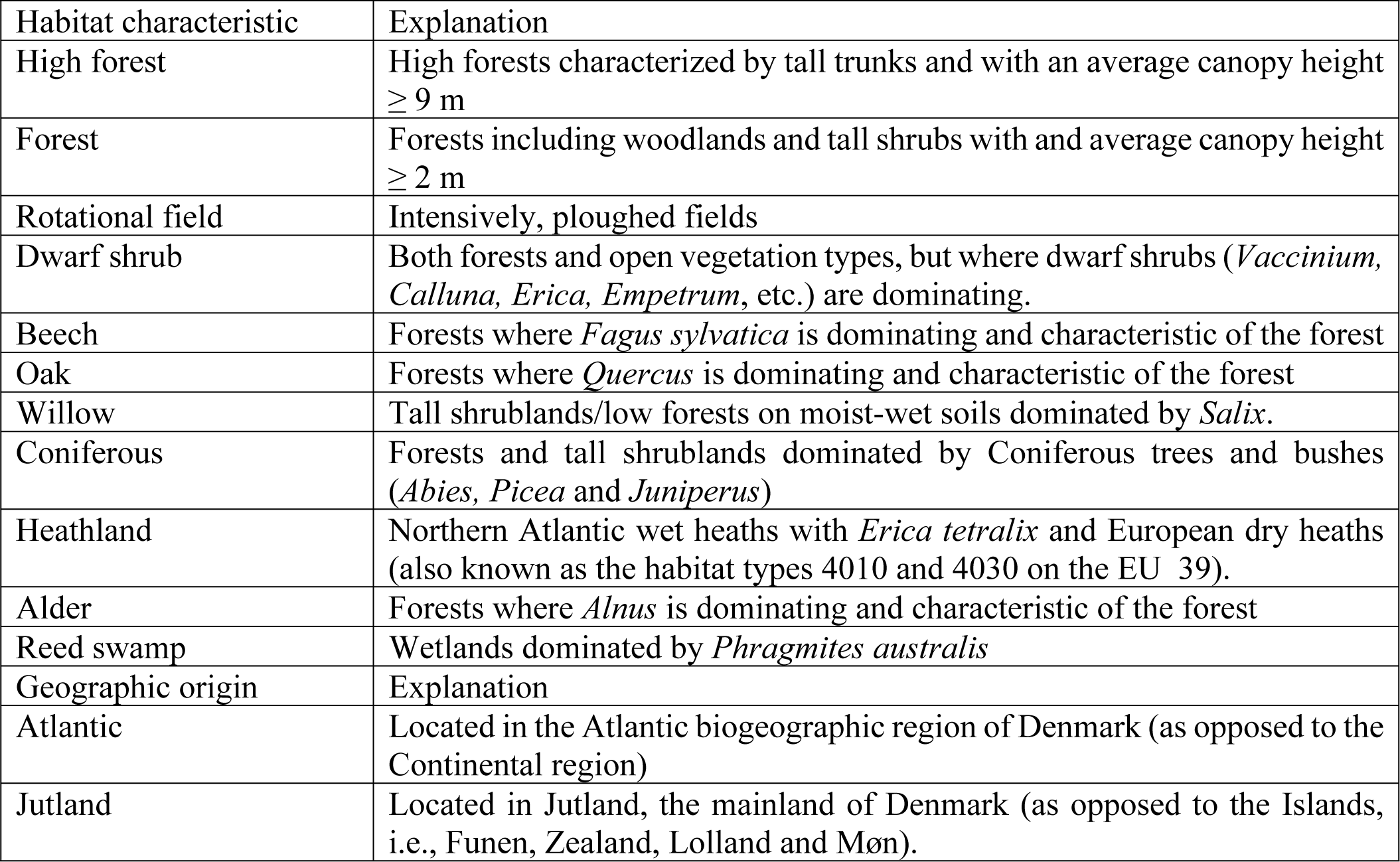
Habitat classes and geographic origins and their description. The classes are binary, i.e., the characteristic is either present or absent at a site.

The habitat classes were modelled using Quadratic Discriminant Analysis using the function qda in package MASS (37) and leave-one-out cross validation was used for model selection and estimation of prediction error. The explanatory variables were the previously mentioned set of NMS axes of the four different OTU-groups (eukaryotes, fungi, plants and insects). Model selection was performed by adding variables that improved the percentage correctly predicted in the cross validation. We used the class proportions for the training set as prior probabilities. As the distribution of the binary characteristics is often skewed with few sites with class membership = 1, percent correctly predicted tends to be high solely because of a high percentage true negatives. Therefore, we also evaluated model performance on the percentage of true positives.

We also investigated the possibility to predict geographic origin defined as a binary response variable, i.e., mainland/islands and Atlantic/Continental biogeographic regions of Denmark, using the above mentioned predictors.

The best model was then selected based on the highest percentage correctly predicted and when that was very similar, we also evaluated the percentage true positives.

For the habitat classes that were characterized by one or a few species, we tested if adding the frequency of annotated sequences improved the model fit. We only used annotated sequences with a 99 % match. Based on our knowledge of the species pool in Denmark, the following frequency of sequences (number of sequences of the target taxon divided by the total number of sequences in the sample) were added as explanatory variables to the best models: *Fagus sylvatica* for beech forest, *Quercus* for oak forest, Pinaceae for coniferous forest, Ericaceae for heathland, *Alnus* for alder swamp and *Phragmites australis* for reed swamp. For willows, the only annotated *Salix* species with a match ≥99% was *Salix herbacea*, which is neither occurring in Denmark, nor indicative of the tall willow scrub here defined as ‘Willow’. However, we also test whether the frequency of sequences of *Salix* improve the model for ‘Willow’.

## Looking for rare and cultivated species

Most plants are rare, i.e., either confined to a rare habitat type or geographically confined and, in the latter case, they may be useful for assessing geographic provenance of a soil sample. Rare plants may occur at single sites in the dataset or at very low frequency of sequences and are therefore not likely to drive geographic patterns in the whole dataset. Instead, we looked for such species as a separate exercise. For this approach, we only used sequences with 100 % GenBank match. We used the nomenclature in Genbank (NCBI taxonomy). In case the sequences matched several species equally well, we used the most frequently referenced species in the database. We checked if the annotated species were rare, i.e., occurring in less than 130 atlas grid cells, corresponding to c. 10 % of the grid cells in the Atlas Flora Danica survey (40), a national mapping of plant species in Denmark in 5 × 5 km grid cells. In addition, we identified 57 plant species in the annotated sequences with 100% match, which are regionally or locally confined in their distributions (see Supplementary materials, Table S1). Cultivated plants, such as common crops, may also help predicting provenance of soil samples and can be compared to the national mapping of fields and crops. For exemplification, we looked for sequences annotated to *Triticum* species, *Brassica napus*, *Beta vulgaris, Secale cereale*, and *Zea mays*.

## Results

Axes from an NMS ordination of OTU communities were good predictors of environmental gradients of fertility, soil moisture, pH and light as represented by EIVs (0.73≤R^2^≤0.88 for the best models, Table 2). NMS axes based on fungal OTUs alone were the best predictors of soil pH and light, whereas eukaryote OTUs best predicted soil moisture and plant OTUs best predicted soil fertility. Although insect OTUs predicted soil moisture and pH well, they were outperformed by fungi and eukaryotes (Table 2).

**Table 2.** Linear model results for predicting environmental gradients. Model R^2^ for each set of predictors based on the different primers and after model selection. Text in bold indicates the best model for each environmental gradient used for prediction.

| EIV | Fungi | Insects | Eukaryotes | Plants |
| --- | --- | --- | --- | --- |
| n | 130 | 128 | 129 | 130 |
| Fertility | 0.81 | 0.56 | 0.80 | **0.83** |
| Soil moisture | 0.82 | 0.82 | **0.86** | 0.79 |
| pH | **0.88** | 0.71 | 0.87 | 0.70 |
| Light | **0.73** | 0.47 | 0.67 | 0.69 |

NMS axes also showed good discriminatory power as predictors of habitat classes in QDA (Table 3). Based on leave-one-out cross validation, most models performed well, with >89% correctly predicted sites. The percentage of true positives was > 50% for all predictions, except Willow, and notably the habitat classes forest, high forest, dwarf shrub, beech, coniferous and reed swamp performed well, i.e. true positives > 67 %. Models did not discriminate well between geographical regions within Denmark and therefore geographic provenance was not predicted. However, we identified 67 plant species in our data set that have geographically distinct distribution patterns (Table S1) and presence of which in a sample may indicate a geographic region. In 95 out of the 130 samples we found sequences of at least one of the 67 regionally or locally distributed species (results not shown).

**Table 3:** QDA model results for predicting habitat classes. Percent correctly predicted binary classes based on the different primers and after variable selection. Percent true positives are reported in parenthesis. For habitats characterized by one or a few species, we tested whether the frequency of sequences would improve model fit; these are indicated by italics. Text in bold indicates the best model selected. When model performances are equal, we opt for the simpler model.

| Binary landscape characteristic | Fungi | Insects | Eukaryotes | Plants | + sequences |
| --- | --- | --- | --- | --- | --- |
| High forest | 0.88 (0.70) | 0.89 (0.70) | 0.87 (0.77) | **0.89 (0.78)** |  |
| Forest | 0.93 (0.91) | 0.85 (0.81) | **0.95 (0.94)** | 0.95 (0.92) |  |
| Rotational field | 0.96 (0.00) | 0.96 (0.00) | 0.98 (0.40) | **0.98 (0.60)** |  |
| Dwarf shrub | **0.91 (0.79)** | 0.79 (0.00) | 0.91 (0.79) | 0.89 (0.71) |  |
| Beech | **0.95 (0.82)** | 0.88 (0.47) | 0.95 (0.81) | 0.94 (0.65) | 0.95 (0.82) |
| Oak | 0.92 (0.00) | 0.91 (0.00) | 0.91 (0.00) | **0.95 (0.64)** | 0.93 (0.45) |
| Willow | 0.92 (0.00) | 0.92 (0.00) | 0.93 (0.00) | **0.95 (0.40)** | 0.91 (0.40) |
| Coniferous | 0.94 (0.11) | 0.93 (0.00) | 0.93 (0.00) | *0.96 (0.44)* | **0.95 (0.67)** |
| Heathland | **0.96 (0.50)** | 0.95 (0.00) | 0.95 (0.00) | 0.95 (0.00) | 0.96 (0.50) |
| Alder | *0.92 (0.53)* | 0.89 (0.27) | 0.91 (0.53) | 0.88 (0.00) | **0.95 (0.60)** |
| Reed swamp | 0.89 (0.61) | **0.90 (0.67)** | 0.89 (0.56) | 0.89 (0.33) | 0.88 (0.17) |

| Binary geographic origin |  |  |  |  |
| --- | --- | --- | --- | --- |
| Atlantic | 0.73 (0.00) | 0.74 (0.00) | 0.73 (0.00) | 0.73 (0.00) |
| Jutland | 0.65 (0.92) | 0.65 (0.81) | 0.66 (0.90) | 0.60 (1.00) |

We present two mock crime scenes as examples of how these predictions can be collated and presented to investigators in a meaningful way (Fig. 2). For mock crime scene 1, the predicted EIVs placed the site at an intermediate position for soil pH and moisture, slightly fertile soil and low light conditions. Considering all four environmental predictions and with the aid of the ellipses in Fig. 1, this indicated mull or moder forest site. The prediction of binary habitat classes showed very low probabilities for open vegetation types (e.g., rotational field and heathland) and wet vegetation (e.g., reed swamp, willow and alder dominated habitat types) and high probabilities for the habitat classes forest, high forest and beech, and some probability for oak and coniferous. The list of plant species and their frequency of sequences in the sample showed that *Fagus sylvatica*, *Tilia cordata* and *Anemone nemorosa* were very frequent and accompanied by other common forest plant species, such as *Melica uniflora* and *Quercus pubescens*. *Q. pubescens* is not found in Denmark and it is probably a related species of *Quercus*, possibly *Q. robus*, which was found at the site in the botanical survey. The geographic distribution of *M. uniflora* in Denmark is limited to the eastern parts (i.e., the continental region, Table S1). The mock crime scene in this case was an old growth beech forest on moder soil in Zealand, Eastern Denmark.

**Fig. 2.**
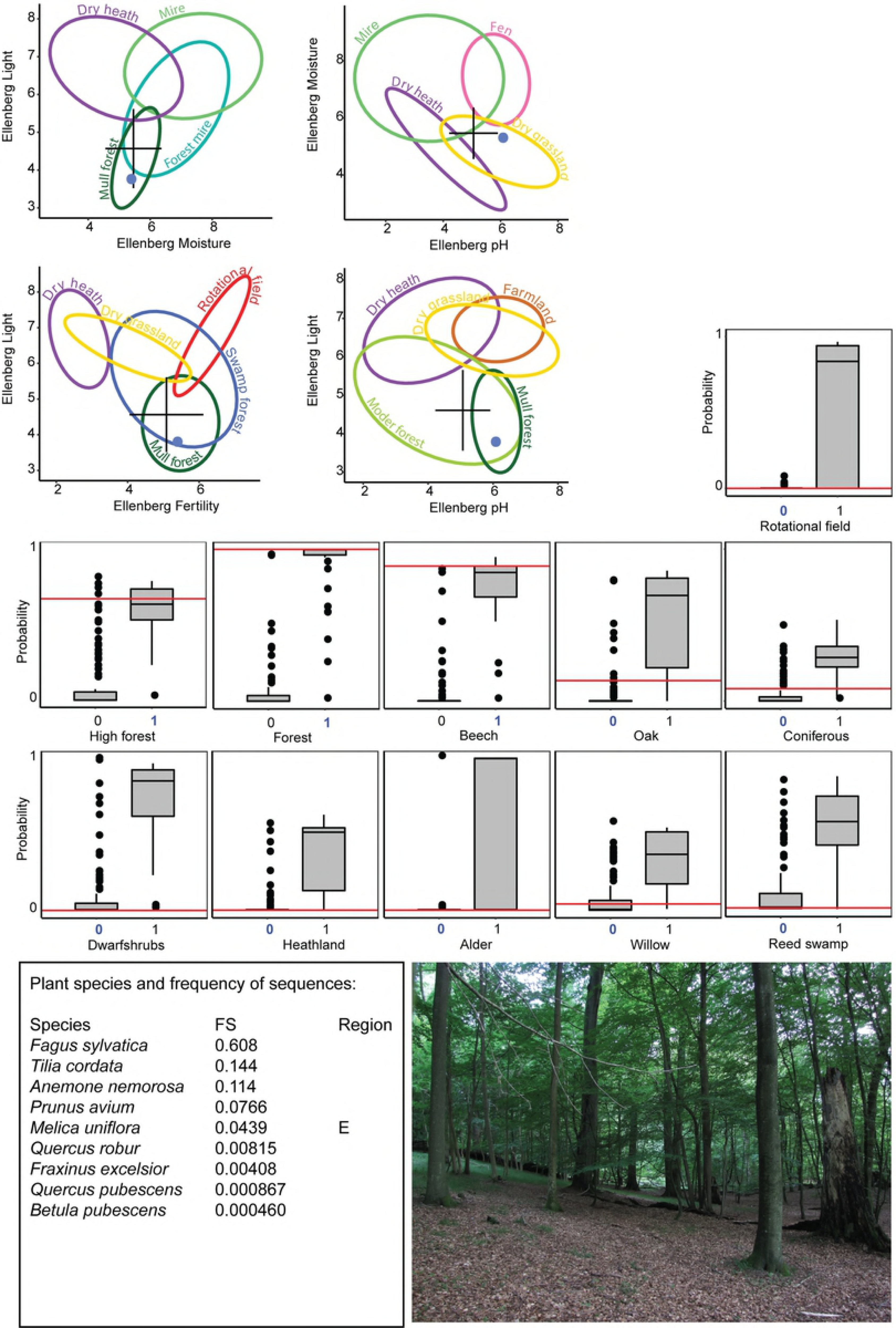
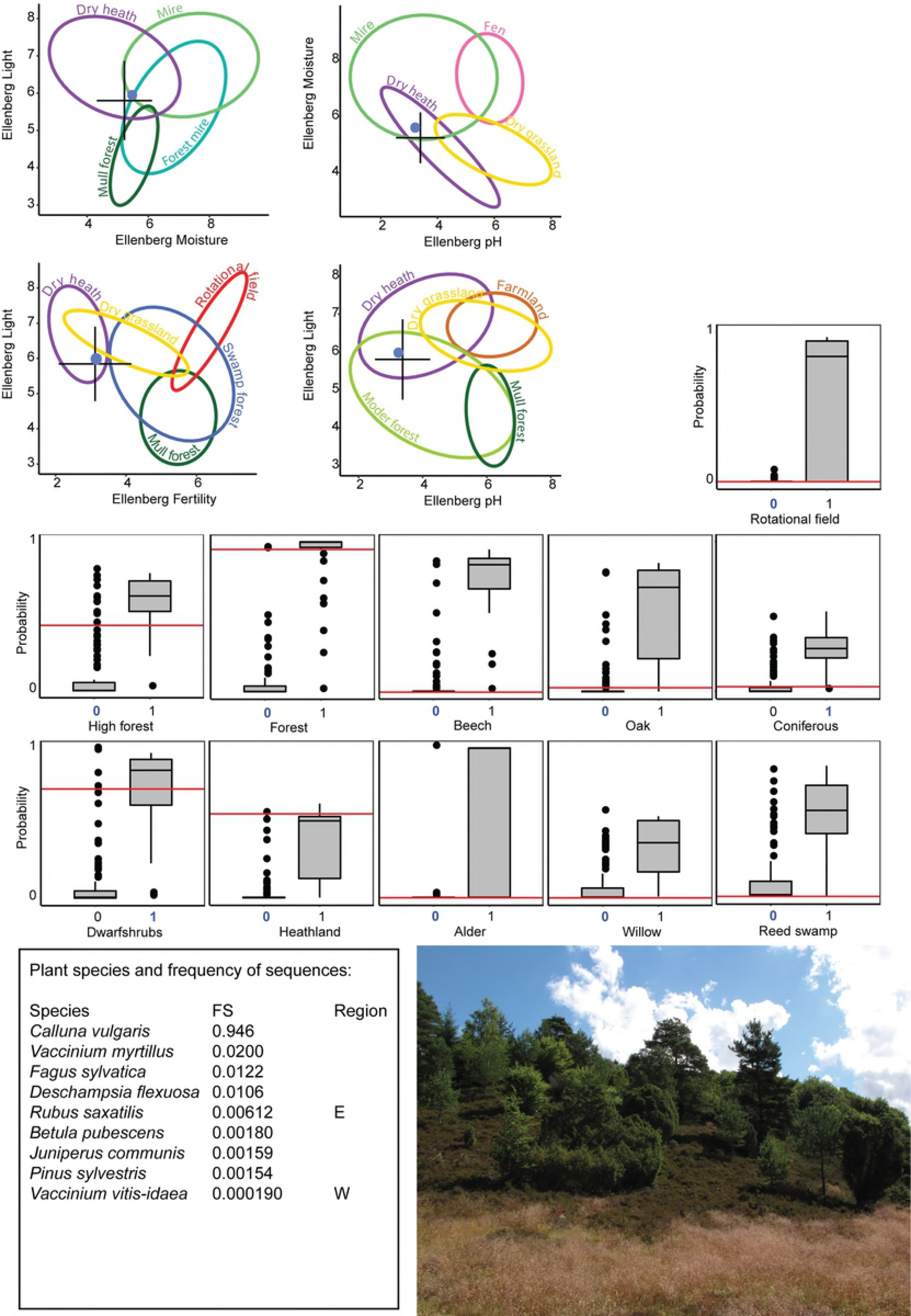
Mock crime scene 1 and 2 and their provenance derived from soil eDNA. Predicted location in environmental space (EIVs) based on the best linear models (top left). The cross marks the predicted EIV value and the length of the arms show the 95% CI. Blue dots show the actual EIVs at the sample site. The predicted probability of binary habitat classes based on QDA are shown as red lines on a box plot of the distribution of predicted values for each characteristic (middle part). A priori classification membership is indicated by blue font color of the 0/1 variable. The probabilities for geographic region are not shown. A list of plant species sequences with 100 % database match, their frequency of sequences (FS) in the sample, and if applicable, the geographic region of their distribution in Denmark is listed bottom left. For evaluation, we provide a picture of the sample location (bottom right).

Predicted EIVs for mock crime scene 2 indicated intermediate light conditions, low nutrient status, low soil pH and relatively low soil moisture, which was within the ellipse of dry heathlands. The binary predictions showed high probability for forest but low probability for high forest. Low probabilities for other forest characteristics, indicates that it is unlikely to be a characteristic beech, oak, or coniferous forest. There were high probabilities for heathland and dwarf shrubs, which corresponds to the annotated species found in the soil sample. The site OTU community has a very high proportion of *Calluna vulgaris* sequences and less of *Vaccinium myrtillus*, *F. sylvatica* and *Deschampsia flexuosa*. *Rubus saxalis* is typical of eastern Danish alkaline forest and the OTU is probably a related species of *Rubus*. Other equally good matches of this sequence in GenBank includes *R. idaeus*, European raspberry, which is much more likely given the other habitat predictions. The recorded species from the botanical field inventory revealed that *R. ideaus* and not *R. saxalis* was found at the site. *Vaccinium vitis-idaea* has a typical westerly distribution. The mock crime scene 2 is a dry heathland with scattered trees *F. sylvatica*, *Q. robur* and *Pinus sylvestris* and *Juniperus communis* shrubs. The site is located in Jutland in Western Denmark.

## Discussion

We investigated the potential for constructing predictive models of environmental properties, habitat classes and geographic origin based on soil eDNA. We found that variation in soil eDNA predicts environmental conditions and habitat classes well, whereas is less well suited for assigning geographical provenance. The latter result corresponds to other ecological studies showing that, within habitat types, geographic variation in species composition is limited within Denmark (e.g., 41). Previous attempts to use eDNA to predict geographic provenance are few and the scale of prediction is important to explore. DNA sequences of fungi in dust have been shown to predict geographic origin with a few hundred km precision at a continental scale (42). For comparison, Denmark extends c. 300 km from east to west. Recently, spectral analysis, another discipline of forensic geoscience, was shown to successfully predict geographic origin at local scales in a cultural landscape (11). It is possible that eDNA also will perform better at predicting local scale provenance within a specific urban landscape, which may be characterized by novel communities and introduced species (43).

Variation in eDNA predicted environmental gradients, i.e., EIVs, in linear models with R^2^ >0.73. We used EIVs for light conditions, soil moisture, nutrient conditions and soil pH (33). Averaging plant indicator values is commonly used in vegetation studies to estimate or assess local conditions (e.g., 34). The validity of plant-based bioindication has been confirmed by direct measurement of the environmental conditions and by plant growth experiments (e.g., 35, 36, 44). EIVs are available only for the Central European flora, however, for application in other parts of the world, EIVs may be replaced by species scores from ordination of large and representative vegetation datasets, which typically reflect major environmental gradients (45).

While habitat types are often described and delimited by distinct characteristics and plant communities (39), in reality habitats are not discrete classes but rather overlapping in species communities and along continuous environmental and successional gradients and occur in patchy mosaics. Performance of our classification models tended to be best for habitats that are relatively well defined and delimited along these gradients, e.g., high forest and forest. On the other hand, we defined heathland broadly to include both wet and dry heathland. Similar communities can be found in mires, grasslands and plantations (see also the overlap in the ellipses in Figure 1) making it difficult to get a high prediction accuracy for heathland.

The two mock crime scenes showed that model predictions for environmental conditions corresponded well with the actual EIV’s at the site. Predictions of habitat classes did not always correspond to the *a priori* classifications at a first look, but this could reflect the continuous nature of biotic gradients more than a model failure. For example, mock crime scene 1 is a *Juniperus communis* formation on heathland (5130 on the Habitats Directive) and therefore not classified as a heathland but as the class coniferous by the definitions used here. However, from the picture of the site is it evident that the high probability of heathland is correct.

As the models use ordination axes from OTU communities, they are probably less likely to be influenced by mistakes and biases in sequencing and free from mistakes that may arise from annotating sequences to species. While botanists and ecologists can interpret a lot of habitat characteristics and environmental conditions from a plant species list, particular care must be taken when that species list arises from DNA sequence matches with sequence data from public databases, as these may be misidentified or in other ways erroneously annotated (e.g., 46). Using plant species for further information requires not only expert knowledge on plant ecology, but also on sequencing, bioinformatics and data quality of reference data. For example, in mock crime scene 2, we found *Rubus saxalis*, stone bramble, in the annotated sequences. This species is rare in Denmark and its typical habitat is forest on alkaline soil. This does not match well with the other habitat classes and the relatively low pH predicted by the statistical models. A few more examples of errors of this type are evident, e.g., low quantities of *Quercus pubescens*, which does not occur in Denmark, but shows up on the annotated sequences in mock crime scene 1 is most likely misidentified *Q. robur* sequences, and in Table S1 *Anthoxanthum aristatum* is most likely the more common *A. odoratum*. In this controlled study, it was easy to re-interpret the OTU assignment as a more likely related species based on botanical expertise. In forensic cases or in the absence of experts, however, care must be taken when interpreting annotated species as there can be mistakes in the database entries or, notably with low frequency of sequences as these could stem from contamination, but also biases in sequencing and mistakes in the reference database. For real case forensic work, sequence annotations (and not merely the OTU composition) to infer ecological provenance, should ideally rely on a dedicated reference database of sequences from relevant regional plants, and only unambiguous species matches should be used in the ecological inference to ensure validity, accuracy and reproducibility.

For further work and application of provenancing soil samples, a national database of soil characteristics including variation in soil eDNA, is essential. For forensic investigations an obvious disadvantage of the present investigation is the lack of urban areas, i.e., parks, rubbish dumps, roadsides etc. among the 130 reference samples. However, the dataset is a good starting point as the sites were stratified according to the spectrum of environmental variation in in Denmark (22).

## Conclusions

Sampling and analyzing soil eDNA allows interpretation of major environmental gradients and habitat classes relevant for both basic and applied ecology, such as forensic ecology. It demonstrates a new application of eDNA and the basic ecological information that can be extracted from eDNA and variation in OTU assemblages. While we demonstrate the potential application of this technique for predicting and interpreting information relevant for forensic investigations, it is also important to note a number of issues that are relevant to explore in the future. As already mentioned, the present dataset (Biowide) was originally designed to explore biodiversity in natural habitats across Denmark, and as such, urban areas are not represented and agricultural fields and other cultural areas are underrepresented. We know little about the seasonal variation in eDNA (but see e.g., 47) and variation in eDNA with soil depth (see e.g., 48). Moreover, forensic soil samples can be minute and contaminated, dried, old and degraded, and we need to explore the model sensitivity and provenancing accuracy of such samples (see also 10).

Here we demonstrate a new application of eDNA and show that the generated results can be used for sample provenancing relevant for forensic investigations. Hence, the eDNA spproach will be a useful investigative tool in crime scene cases without the need for the strict and validated procedures necessary for comparisons that have to be used as evidence in court. Given the generic structure of the eDNA approach we expect it to have global relevance by being applicable in a number of countries outside Denmark.

## Acknowledgements

We consider this a contribution to a national research project to develop soil forensic methods for Denmark supported by Innovation Fund Denmark Grand Solutions (grant no. 6151-00002B, https://innovationsfonden.dk/, awarded to AJH) based on data collected in Biowide, a nationwide survey of biodiversity in Denmark supported by the VILLUM foundation (grant no. VKR-023343, https://veluxfoundations.dk/, awarded to RE).

## Supporting information

S1: List of rare species and plant species with geographically limited distribution

Author contributions: AKB, RE, TGF and HHB contributed to the field inventory of biodiversity in Biowide. CF, TGF, AJH and RE conceived the idea. TGF lead the laboratory and bioinformatics analyses. CF, JME and RE analyzed the data. HHB and AJH contributed with botanical and forensic expertise, respectively. CF lead the writing and all other contributed.

